# Comparative Molecular Genomic Analyses of a Spontaneous Rhesus Macaque Model of Mismatch Repair-Deficient Colorectal Cancer

**DOI:** 10.1101/2021.08.19.456691

**Authors:** Nejla Ozirmak Lermi, Stanton B. Gray, Charles M. Bowen, Laura Reyes-Uribe, Beth K. Dray, Nan Deng, R. Alan Harris, Muthuswamy Raveendran, Fernando Benavides, Carolyn L. Hodo, Melissa W. Taggart, Karen Colbert Maresso, Krishna M. Sinha, Jeffrey Rogers, Eduardo Vilar

## Abstract

Colorectal cancer (CRC) remains the third most common cancer in the US with 15% of cases displaying Microsatellite Instability (MSI) secondary to Lynch Syndrome (LS) or somatic hypermethylation of the *MLH1* promoter. A cohort of rhesus macaques from our institution developed spontaneous mismatch repair deficient (MMRd) CRC with a notable fraction harboring a pathogenic germline mutation in *MLH1* (c.1029C<G, p.Tyr343Ter). Our study incorporated a detailed molecular characterization of rhesus CRC for cross-comparison with human MMRd CRC. We performed PCR-based MSI testing, transcriptomic analysis, and reduced-representation bisulfite sequencing (RRBS) of rhesus CRC (n=41 samples) using next-generation sequencing (NGS). Systems biology pipelines were used for gene set enrichment analysis (GSEA) for pathway discovery, consensus molecular subtyping (CMS), and somatic mutation profiling. Overall, the majority of rhesus tumors displayed high levels of MSI (MSI-high) and differential gene expression profiles that were consistent with known deregulated pathways in human CRC. DNA methylation analysis exposed differentially methylated patterns among MSI-H, MSI-L (MSI-low)/MSS (MS-stable) and LS tumors with *MLH1* predominantly inactivated among sporadic MSI-H CRCs. The findings from this study support the use of rhesus macaques as the preferred animal model to study carcinogenesis, develop immunotherapies and vaccines, and implement chemoprevention approaches pertinent to sporadic MSI-H and LS CRC in humans.

## Introduction

Colorectal cancer (CRC) remains the third leading cause of cancer-related deaths affecting both men and women (1). Approximately 15% of CRC cases display microsatellite instability (MSI) secondary to a defective mismatch repair (MMRd) system that is recognized as a major carcinogenic pathway for CRC development. MMRd arises as a result of either (1) an inherited germline mutation in one of four genes (*MLH1, MSH2, MSH6* and *PMS2*) constituting the MMR system followed by an acquired second-hit in the wild-type allele of the same gene in colonic mucosa cells (i.e., Lynch syndrome) or (2) somatic inactivation of the *MLH1* gene (i.e., MSI sporadic CRC).

A better understanding of colorectal neoplasia arising in the setting of MSI/MMRd is urgently needed to tailor the use of early detection, prevention, and treatment interventions in this subset of CRC, including established immunotherapies and the development of novel immuno-preventive regimens. Such interventions are particularly needed for those with Lynch syndrome, as they are at the highest risk of CRC as well as a range of other cancers. Unfortunately, no concrete model with higher translational value exists to study the nuanced carcinogenesis of MMRd CRC, which is a critical barrier for studying this subset of CRC and, consequently, to making advances in its detection, prevention, and treatment.

Presently, *in vitro* and *ex vivo* models, such as cell lines and organoids respectively, are commonly used to study CRC; however, the intrinsic nature of these models lack cellular heterogeneity and fail to recapitulate the tumor microenvironment (TME) observed *in-vivo* (2). To combat the limitations of *in-vitro/ex-vivo* cultures, mouse models (*Mus musculus*) have been leveraged to study CRC prevention, initiation, and progression. Although murine models of genetic inactivation of MMR genes exist, these models drastically diverge from the human LS (MMRd) phenotype. For example, murine models with constitutional homozygous MMR gene inactivation have high rates of lymphoma formation, limiting the efficacy of these models. In an effort to circumvent this challenge, investigators have employed tissue-specific Cre recombinase-based inactivation of MMR genes; however, these mice predominantly develop tumors in the small intestine (as opposed to the large intestine in humans) (3). These limitations of cellular cultures and murine models warrant the need for better model systems to elucidate the intrinsic and extrinsic factors of MMRd carcinogenesis to help improve clinical outcomes for both LS and MSI sporadic patients.

Given the anatomic and physiologic similarities and genomic homology between non-human primates (NHPs) and humans, researchers have used several species of NHPs to develop therapies and vaccines to treat and eradicate human disease (4, 5). The rhesus macaque (*Macaca mulatta*), which shares 97.5% DNA sequence identity with humans in exons of protein-coding genes as well as close similarity in patterns of gene expression, has been an invaluable animal model for studying human pathophysiology (6, 7). Studies have shown that rhesus launch parallel immune responses and display analogous pathologies to humans, thus making them ideal animal models suited for clinical translation of basic and pre-clinical findings compared to other model organisms (8-11).

A cohort of specific pathogen free (SPF), Indian-origin rhesus macaques bred at The University of Texas MD Anderson Cancer Center (MDACC) Michale E. Keeling Center for Comparative Medicine and Research (KCCMR) spontaneously develops MSI/MMRd CRC, including a subset of animals harboring a pathogenic germline mutation in *MLH1* (c.1029C<G, p.Tyr343Ter). This spontaneous mutation manifests into clinical and pathological features similar to human LS, which suggests that these rhesus macaques may be a superior model organism for studying MMRd CRC (10, 12).

This study characterized the genomic features of colorectal tumors in the KCCMR rhesus cohort using microsatellite marker testing, whole transcriptomics, and epigenomics coupled with systems biology tools, as illustrated in **Figure 1**. Additionally, we cross-compared the current subtypes of CRC in humans with the rhesus model to evaluate the utility of rhesus for studying early cancer development, treatment modalities, and prevention approaches in hereditary and sporadic CRC.

**Figure 1.**
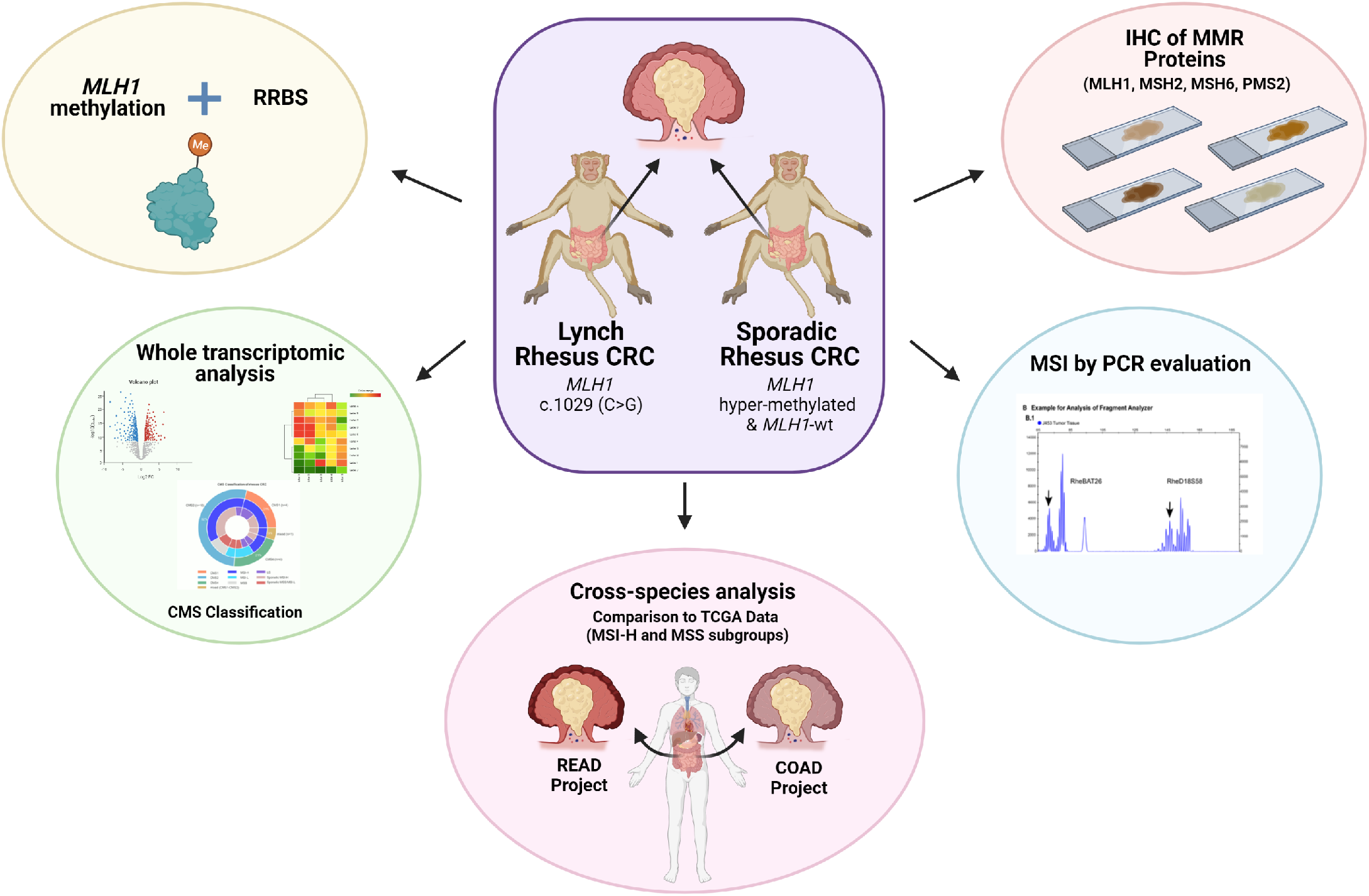
Schematic outline of the experimental design. Sporadic and rhesus Lynch (heterozygous *MLH1* nonsense mutation, c.1029, C>G) animals bred and housed at UTMDACC KCCMR were used to genomically characterize colorectal tumors using an in-house MSI panel, IHC of MMR proteins, epigenetic evaluation, whole transcriptomic analysis, and CMS classification. These analyses establish the framework for utilizing rhesus as a surrogate to study MMRd CRC. UTMDACC KCCMR, University of Texas MD Anderson Cancer Center Michale E. Keeling Center for Comparative Medicine and Research; MSI, microsatellite instability; MMRd, mismatch-repair deficiency; CMS, consensus molecular subtype; CRC, colorectal cancer.

## Results

### Clinical characteristics of colorectal tumors in rhesus

We identified a total of 41 animals diagnosed with CRC at the time of necropsy. All tumors were located in the right side of the colon (20 in the ascending colon, 16 in the ileocecal valve, and 4 in the cecum) with the exception of one jejunal tumor. The mean age at death was 19.3 years (range: 9 and 27 years, **Figure 2A**) and 80% of animals were female (**Figure 2B**), consistent with overall population demographics of approximately 80% females from which the CRC animals were drawn. The average age at death was younger among the LS macaques than among the sporadic MSI macaques, but the difference is not statistically significant (17.75 vs 19.75 years, *P*-value=0.3, **Figure S1**).

**Figure 2.**
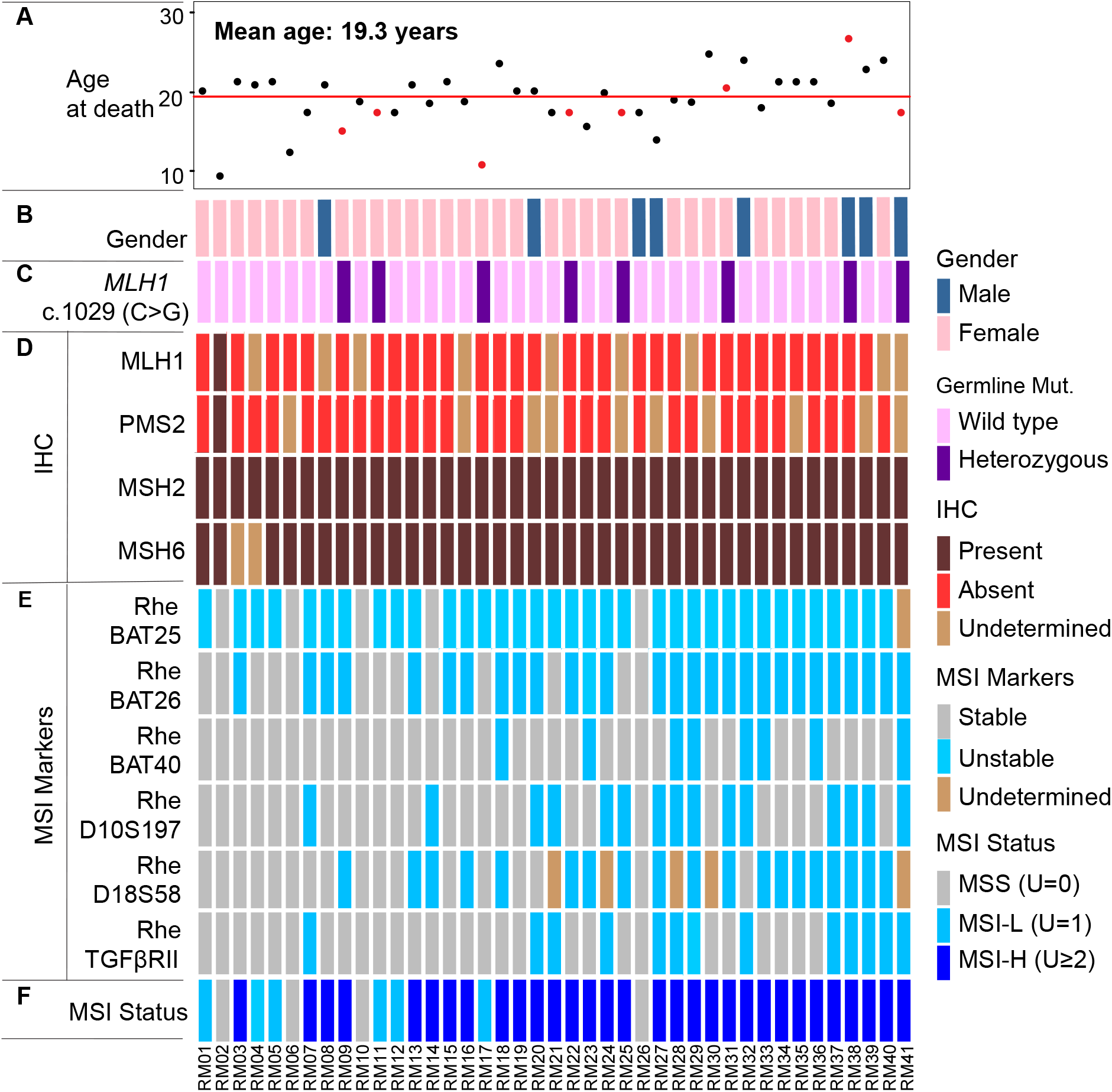
Clinical, pathological, and molecular characteristics of the Rhesus cohort. (**A**) Animal ages at the time of diagnosis of CRC and subsequent euthanasia. The average age at death for the rhesus CRC cohort was 19.3 years. Red dots denote the age of animals with *MLH1* germline mutation; (**B**) Gender of KCCMR rhesus cohort. The majority of animals in this cohort were female; (**C**) Lynch syndrome *MLH1* germline mutation status. Out of forty-one animals, eight (20%) carried a heterozygous *MLH1* nonsense mutation (c.1029, C>G); (**D**) IHC assessment of rhesus CRC. The majority of tumor samples of rhesus CRC displayed loss of MLH1 and PMS2; (**E**) MSI testing of rhesus tumors. A newly designed MSI testing panel for rhesus CRC included six markers (RheBAT25, RheBAT26, RheBAT40, RheD10S197, RheD18S58, and RheTGFβRII) that were orthologs of commonly tested MSI loci in human tumors (BAT25, BAT26, BAT40, D10S197, D18S58, and TGFBRII). Overall, RheBAT25, RheBAT26, and RheD18S58 MSI markers were the most mutable MSI markers in rhesus CRC; (**F**) Summary of MSI status of rhesus tumors. Rhesus CRC were predominantly MSI-H (75%), and only six tumors (15%) were MSI-L, and four (10%) MSS.

### Germline Genetics

We detected the presence of a previously described heterozygous germline stop codon mutation in exon 11 of *MLH1* (c.1029C>G; p.Tyr343Ter, **Figure 2C, Figure S2 and Table S1**) in 8 animals (∼20%) from KCCMR (10), thus confirming the presence of a causative pathogenic mutation of Lynch syndrome in humans (herein referred to as rhesus Lynch) (12). The remaining 33 animals (80%) had the wild-type germline variant of *MLH1* (herein referred to as rhesus sporadic, **Figure 2C**).

### Immunohistochemistry (IHC) staining displayed widespread loss of expression in MLH1 and PMS2 in rhesus CRC

Of the rhesus CRCs with IHC data (n=37), 36 samples (97%) had loss of MLH1 and/or PMS2 protein expression. Only one animal (∼3%) retained the expression of the MLH1-PMS2 heterodimer. This same animal also displayed complete stability of the MSI markers, thus being MSS, and therefore, was considered MMR proficient. We subsequently used this animal as a control for all further genomic analyses (**Figure 2D**).

### Assessment of MSI in rhesus CRC

We developed an MSI testing panel for rhesus CRC including orthologs of the most frequently used microsatellite markers in human CRC: BAT25, BAT26, BAT40, D10S197, D18S58, D2S123, D17S250, D5S346, β-catenin, and TGFβRII. Rhesus orthologs of D2S123, D17S250, and D5S346 markers did not contain adequate nucleotide repeats suitable to assess the presence of MSI. Hence, we excluded these markers from the rhesus MSI testing panel. Furthermore, the rhesus ortholog of BAT25 was not sensitive enough to determine MSI in rhesus CRC due to the interruption of the microsatellite by a nucleotide. Therefore, we substituted it with a novel MSI marker— c-kitRheBAT25—identified through screening the whole sequence of the *c-kit* gene for an uninterrupted repeat region. Overall, the rhesus CRC MSI testing panel included 6 markers: 4 mononucleotide (c-kitRheBAT25, RheBAT26, RheBAT40, RheTGFβRII) and 2 dinucleotide (RheD18S58, RheD10S197) markers (**Table S1**). This panel offers an assessment of the functionality of the MMR system in these rhesus macaques.

With the newly designed rhesus MSI panel, we performed MSI testing of tumors from the entire KCCMR cohort and used matched normal samples as a genomic reference (n=41). c-kitRheBAT25, RheBAT26, and RheD18S58 markers were the most sensitive (**Figure 2E**). We validated the calls made in RheBAT26 and RheD18S58 using an alternative technique based on fragment analysis (**Figure S3**). We classified rhesus tumors into three categories—MSI-H, MSI-L, MSS—by counting the number of unstable markers in each tumor and abided by classical NCI recommendations (13). Thirty-one samples were MSI-H (76%, herein referred to as rhesus sporadic MSI), six were MSI-L (15%), and four were MSS (10%) (**Figure 2F**).

### DNA methylation was responsible for developing CRC in the rhesus

As seen in human MSI CRC, the phenotype of rhesus MSI-H CRC determined from MSI testing and transcriptomic profiling suggests a vast majority of rhesus CRC may involve an epigenetic event. To determine the epigenetic contribution to rhesus CRC, we analyzed the global DNA methylation patterns in tumor and normal samples.

Unsupervised principal component analysis (PCA) of reduced-representation bisulfite sequencing (RRBS) data revealed clear clustering of MSI-H, MSI-L/MSS, and Lynch syndrome tumors, as well as normal mucosa (**Figure 3A**). Hierarchical clustering of DNA methylation profiles using Pearson’s correlation distance displayed a clear separation between rhesus tumor and matched normal tissue samples. Rhesus MSI-H tumor tissue samples clustered together with rhesus LS and separated from normal and rhesus MSS/MSI-L CRC (**Figure 3B**). Significant differentially methylated regions (DMRs) between rhesus normal and tumor tissue samples using a FDR of 5% involved the following genes: *TOP1, PCGF3*, and *FAM76B* (hypermethylated), and *ALKBH5, GAS8*, and *MME* (hypomethylated, **Figure 3C**).

**Figure 3.**
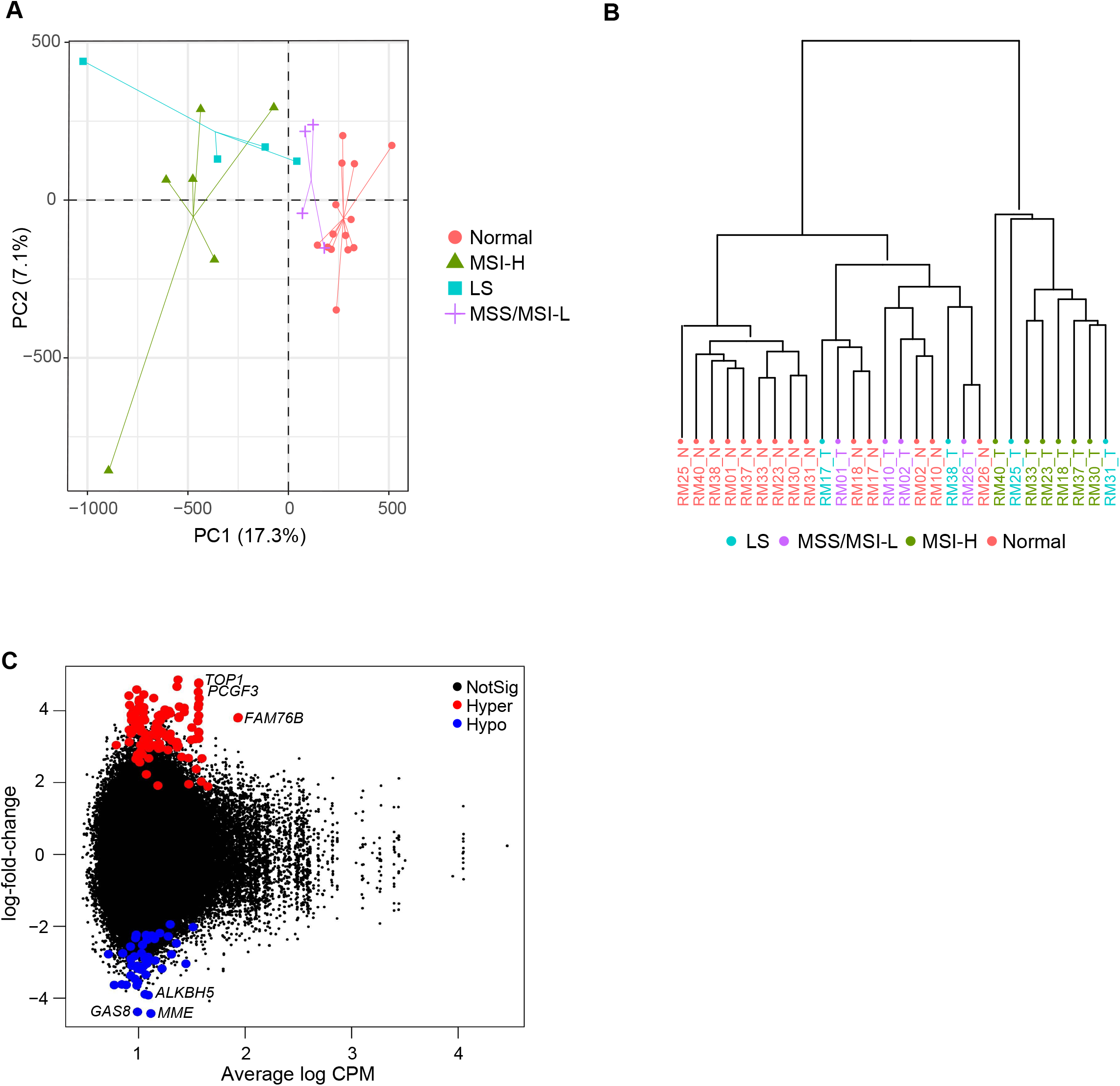
Methylation analysis of rhesus CRC. (**A**) PCA of DNA methylation in rhesus specimens characterizing the trends exhibited by the differentially methylated region profiles of sporadic MSI-H (green triangle), sporadic MSS and MSI-L (purple plus), Lynch syndrome (blue square), and normal tissue (red circle) samples. Each shape represents a tissue sample type. Each group clustered separately; (**B**) Hierarchical clustering of DNA methylation profiles assessed by CpG methylation using Pearson’s correlation. Distance displays the relationship between rhesus tumors and matched normal tissue samples with parameters set as distance method: “correlation”, clustering method: “ward”; (**C**) Significant differentially methylated regions (DMRs) of rhesus normal and tumor samples at FDR of 5%. *TOP1, PCGF3* and *FAM76B* were some of the hyper-methylated genes, and *GAS8, ALKBH5* and *MME* were hypo-methylated genes in rhesus CRC.

Lastly, we performed a dedicated methylation analysis of the *MLH1* promoter using a methyl NGS panel. Locations of CpG regions were shown from the transcription start site of the *MLH1* gene. Overall, thirteen CpG regions were significantly methylated in rhesus sporadic MSI-H tumor samples (*P-*value<0.05) compared to adjacent normal mucosa. The majority of methylated CpG regions were located within exon 1. There were no significant methylation differences between other tumor sub-groups and normal tissue samples (**Figure S4)**; however, there was a clear trend of higher levels of *MLH1* promoter methylation among rhesus sporadic MSI-H compared to MSS tumors as well as a notorious absence of *MLH1* methylation in the only LS tumor tested, which is consistent with human CRC biology.

### Gene expression patterns displayed differences between rhesus colorectal tumor and adjacent normal mucosa

Then, we performed whole transcriptome sequencing in 21 colorectal tumors and twenty matched normal mucosa samples. We had to exclude two tumors and four normal samples from downstream analysis due to low mapping efficiency. Unsupervised principal component analysis (PCA) of RNAseq data showed a clear separation of tumor and normal samples. However, samples from rhesus LS, rhesus sporadic MSI-H, and rhesus MSS/MSI-L clustered together without clear separation (**Figure 4A**). Additionally, to further characterize the rhesus LS animal model for studying human MSI-H colorectal cancer, we compared the similarity between rhesus LS tumor samples and human MSI-H and MSS colorectal tumors samples. The differential gene expression between The Cancer Genome Atlas (TCGA) colorectal adenocarcinoma (COAD and READ, respectively) MSI-H tumor samples (n=96) vs. the COADREAD MSS tumor samples group (n=440) was analyzed by edgeR package. One hundred and one orthologous genes demonstrated statistically significant (BH-adjusted *P*-value < 0.05) changes in the expression level by at least two-fold (log2FC≥1). Then we compared the spearman correlation between the rhesus Lynch tumor samples (n=21) and COADREAD MSI-H and MSS samples, while we used COADREAD normal (n=54) and rhesus normal samples (n=20) as control of species distance. The rhesus Lynch tumor samples have a larger correlation with COADREAD MSI-H tumor samples (0.82) than that with COADREAD MSS samples (0.68) and normal samples (0.64, **Figure 4B**). This suggests that our analysis has sufficient resolution to compare different tumor tissue similarities.

**Figure 4.**
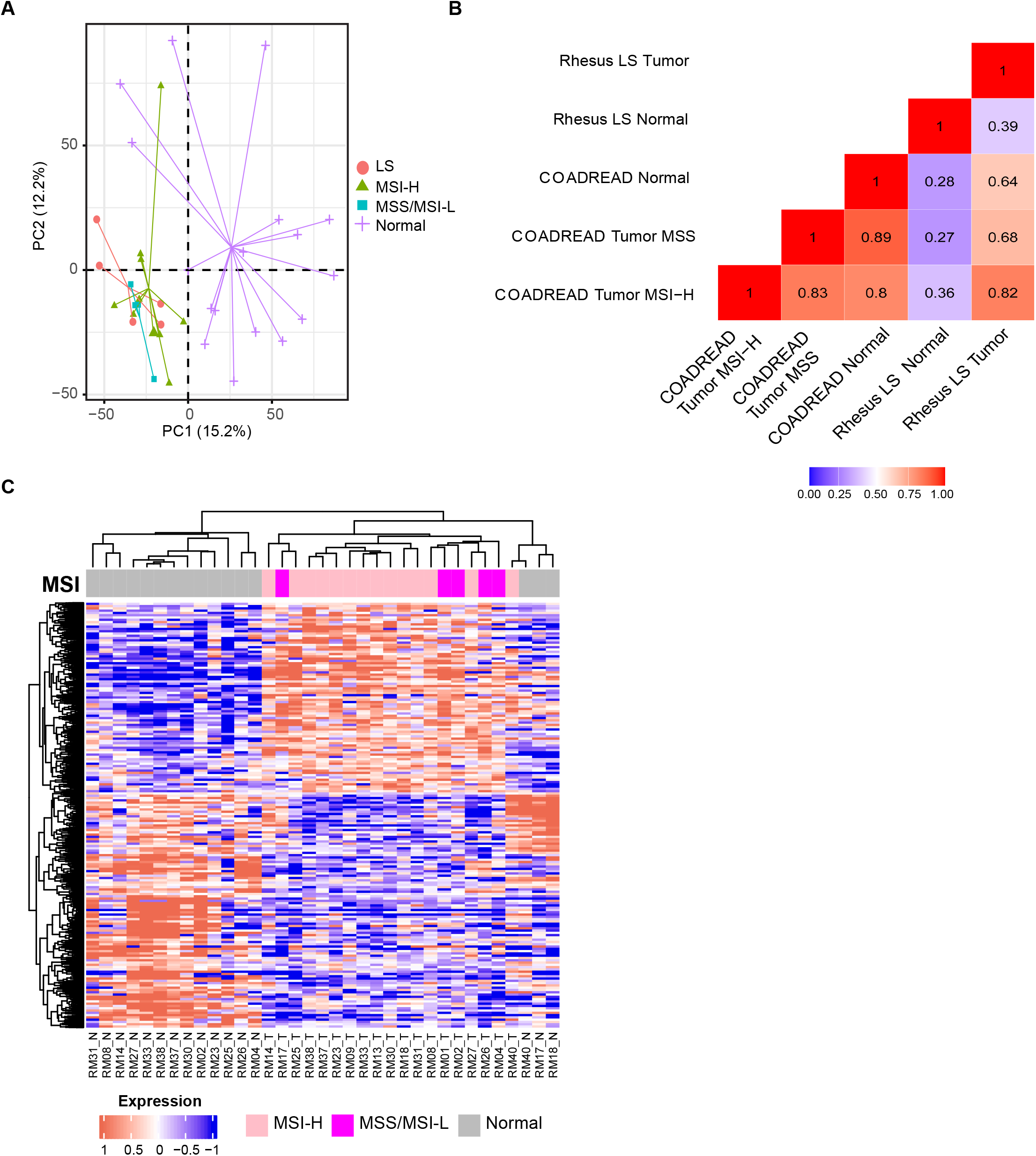
Transcriptomic analysis of rhesus CRC. (**A**) Principal component analysis (PCA) of rhesus CRC showed the trends exhibited by the expression profiles of sporadic MSI-H samples (green triangles), sporadic MSS and MSI-L (blue squares), Lynch syndrome (red circles), and normal tissue (purple plus signs). Normal tissue samples clustered separately from tumor tissue samples; (**B**) Pearson’s correlation coefficient of mean expression levels across 101 significant genes from COADREAD MSI-H tumor samples, COADREAD MSS tumor samples, COADREAD normal tissue samples, rhesus LS tumor samples, and rhesus normal tissue samples; (**C**) Significant differentially expressed genes (DEGs) between tumor and normal tissue samples. DEGs were found based on BH-adjusted *P-*value≤0.05 between rhesus colorectal normal and tumor. Pearson’s correlation was used to perform hierarchical clustering between rhesus tumor and normal tissue samples. Columns represent samples, and rows represent statistically significant differentially expressed genes. Gray color represents normal, pink MSI-H, and magenta MSS and MSI-L tissue samples.

We then determined significantly differentially expressed genes (DEGs) between rhesus normal and tumor by setting a Benjamini-Hochberg (BH)-adjusted *P-*value≤0.05 and log2 fold change±1. We annotated genes using human orthologs (**Figure S5A**). Unsupervised hierarchical clustering using DEGs demonstrated that rhesus tumor tissue samples clustered separately from normal tissue samples, and rhesus MSS/MSI-L CRC were separated from MSI-H CRC samples. Notably, animal RM17 displayed a MSS phenotype despite carrying the *MLH1* germline mutation and clustered with the MSI-H group as opposed to the LS cohort (**Figure 4D**). Using the total RNAseq data, we sought to validate the expression of MMR genes using the counts of reads in tumors and matched normal samples. *MLH1* read counts in MSI-High CRC samples were significantly decreased compared to normal tissue samples (*P-*value<0.0001). As expected, animal RM02 with a MSS tumor showed more *MLH1* read counts in tumor than matched normal (**Figure S5B**). *MSH6* gene read counts in MSI-H CRC samples were significantly more abundant than matched-normal samples (*P-*value<0.001). Differences of *MSH2* and *PMS2* gene read counts between rhesus tumor and normal tissue samples were not significant.

We performed gene set enrichment analysis (GSEA) to discover relevant pathways in colorectal carcinogenesis of MSI-H and MSI-L/MSS rhesus CRC using the ESTIMATE algorithm, which assesses immune and stromal cell admixtures in tumors, canonical, immune, and metabolic pathways (**Figure 5A-C**) (14, 15). When compared with normal tissue samples, the top observed pathways enriched in MSI-H tumor samples involved in cell cycle regulation, crypt base dynamics, and integrin signaling. Conversely, metabolic pathways in MSI-H samples were downregulated compared to normal tissue (**Figure 5A**). A similar trend was observed for MSS/MSI-L tumor samples compared to normal (**Figure 5B**). Lastly, comparing the significant pathways between MSS/MSI-L and MSI-H, we observed an upregulation of key pathways involved in cell cycle regulation and MYC targeting in the MSI-H group (**Figure 5C**).

**Figure 5.**
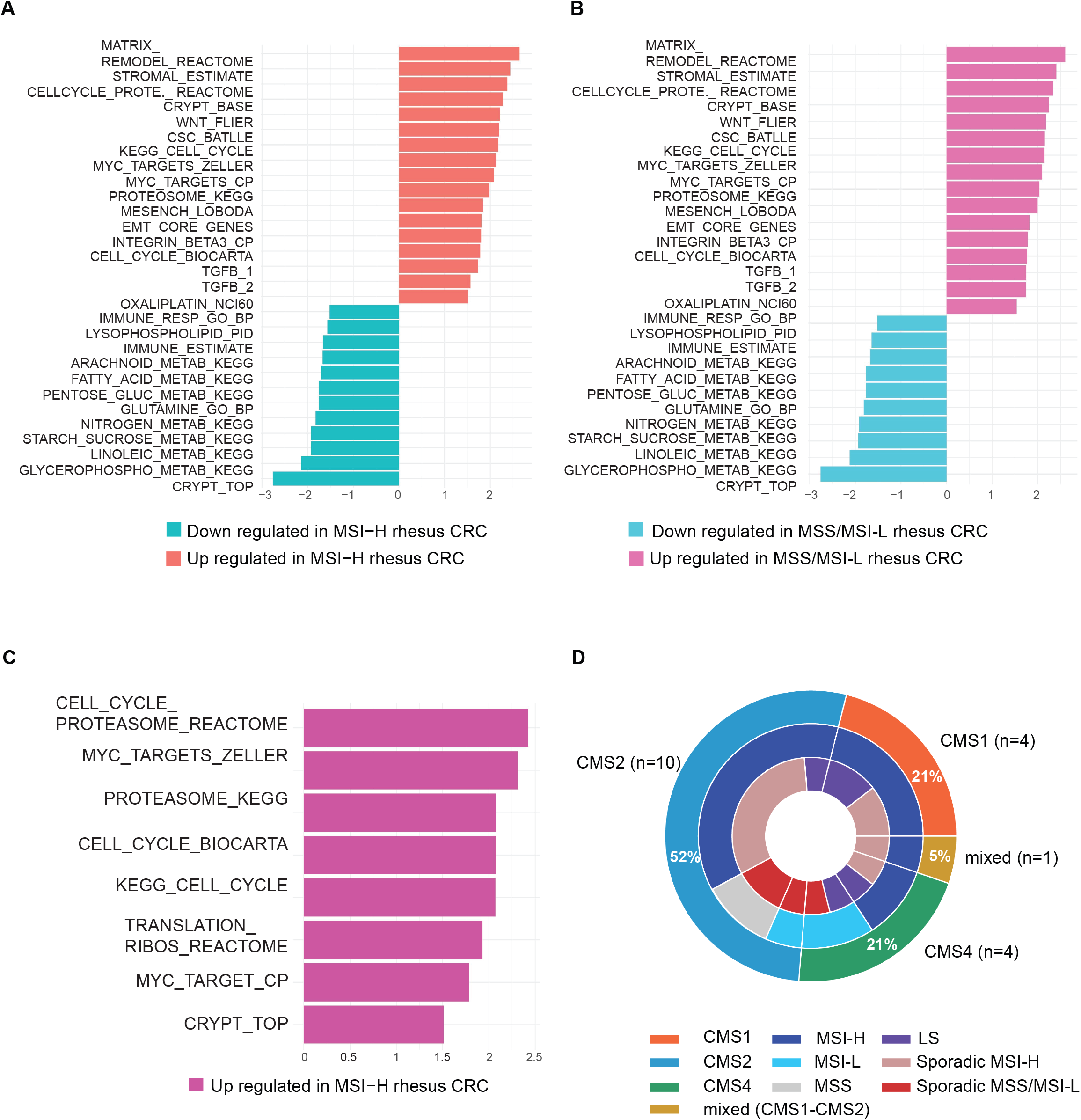
Gene set enrichment analysis in rhesus CRC. (**A-C**) Gene expression pathways are significantly deregulated in rhesus CRC. Pathways relevant to CRC biology are highlighted. BH-adjusted *P-*value≤0.05 was set as a threshold for analysis; (**D**) CMS classification of rhesus CRC. The outer ring of circos plot represents CMS subtypes present in rhesus CRC with 52% of samples (n=10) classifying as CMS2. The middle ring represents the MSI status of samples, and the inner ring indicates clinical categories of samples.

### CMS classification categorized rhesus CRC samples mainly as CMS2

We assigned a consensus molecular subtype (CMS) status to each tumor sample based on the nearest CMS probability (**Table S3**). Overall, 52% (n=10) of tumors were classified as CMS2, which corresponds to the canonical pathways of colorectal carcinogenesis; 21% (n=4) were CMS1, which progresses through MSI and immune pathways; and 21% (n=4) were CMS4, which develops through mesenchymal pathways. Only one tumor displayed mixed features (CMS1-CMS2) of a transition phenotype (**Figures 5D**).

### Rhesus CRC causes mutations in commonly mutated CRC genes

We examined somatic variants of rhesus CRC using total RNAseq data. Our data indicated that the mutation rate of rhesus CRC is relatively high in all tested samples (**Figure S6A**). Commonly altered genes in human CRC were also mutated in rhesus such as *APC, ARID1A, TGBRII, TP53, CTNNB1, PIK3CA, KRAS* (**Figure S6B**). Substitutions of cytosine to thymine were the most abundant in somatic variants of rhesus CRC (**Figure S6C**). Due to the close relation found in humans between MSI-H status and *BRAF* mutations, we performed Sanger sequencing to assess the mutational status of the *BRAF* mutation hotspot *V600E* in rhesus CRC. While we did not detect *BRAF V600E* mutations among rhesus tumors, we did observe different types of *BRAF* somatic variants including missense, nonsense, in-frame, and frameshift deletions (**Figure S7**).

## Discussion

Although cell cultures, organoids, and murine animal models are the most frequently used models in CRC research, these systems fail to recapitulate the phenotypic features of MMRd CRC, which limits clinical translation to humans. To overcome the differences between humans and research models, investigators have turned to NHPs due to their high degree of genomic and physiologic similarity to humans, including natural inter-individual genetic variation. Previous reports have proven rhesus macaques to serve as a durable and clinically-relevant animal model to study many infectious diseases and cancers (9, 10, 16, 17). In this study, our results from MSI testing, IHC, gene expression patterns, systems biology, somatic variant calling, and DNA methylation of colon tissue samples from the KCCMR cohort demonstrated that rhesus macaques develop CRC phenotypes analogous to MSI CRCs, including LS patients. These finding indicate that rhesus macaques may serve as an optimal animal model for studying MMRd CRC and addressing the shortcomings of previously-established model systems.

To characterize the rhesus macaque as a surrogate for studies of MMRd, we investigated the MSI status of 6 markers across 41 unique rhesus tumors using a newly designed, in-house MSI panel for rhesus CRC. Our study results indicated that 76% of rhesus CRC from the KCCMR cohort had a MSI-H phenotype, which warrants the use of rhesus as an optimal system to study MSI-H carcinogenesis. Many rhesus tumors lost expression of MLH1 and PMS2 proteins, but retained the expression of MSH2 and MSH6, as confirmed by IHC analysis. The *MLH1* germline stop codon mutation (c.1029C>G, p.Tyr343Ter), previously reported as a likely pathogenic variant in human LS (National Center for Biotechnology Information), was present in 8 (19.5%) rhesus macaques, while the majority (80.5%) were wild-type for this variant.

The DNA methylation analysis of rhesus CRC in this study suggests that epigenetics plays a pivotal role in rhesus CRC development. DNA methylation status of rhesus CRC using FFPE tissue samples from colon tumor and adjacent normal tissue samples indicated clear segregation of methylation patterns between tumor/normal matches. Furthermore, based on analysis of the RRBS data, DNA methylation appears to play a major role as a driver of rhesus MSI CRC. Interestingly, although human CRC typically displays DNA methylation in the promoter region of the *MLH1* gene, methylation of rhesus CRC predominantly occurred in the exon1 region of *MLH1*.

Despite prior reports of tissue-specific transcriptome analysis of fresh frozen tissues from rhesus macaques, to date, no study has analyzed the transcriptomic profile of colonic tissue from Indian origin rhesus macaques (18). Therefore, our study is the first transcriptomic analysis of matched tumor and normal colon samples in rhesus macaques, which provides essential information for the field of MMRd-related research. Our transcriptomic data of rhesus CRC from FFPE tumor tissue displayed gene expression differences between rhesus tumor and normal tissue samples, and when compared to human TCGA MSI/MSS CRC data, rhesus MSI-H tumors were more similar to human MSI-H expression patterns than were human MSS tumors. These findings of transcriptomic homology between humans and rhesus support utilizing rhesus LS to study the carcinogenesis of MMRd CRC.

To confirm the biological relevance of the rhesus macaque as an animal model, we performed CMS classification and GSEA to ascertain the molecular features of rhesus MSI CRCs, including LS CRCs. Rhesus CRC from predominantly sporadic MSI-H and sporadic MSS/MSI-L mainly associated with CMS2—the canonical subtype—which corresponds to SCNA high and WNT/MYC activation (14). However, rhesus LS tumors primarily associated with CMS1 (MSI-Immune), which aligns with previous studies from our group and encompasses MSI, CpG Island Methylator Phenotype (CIMP) high, hypermutation, immune infiltration, and worse overall survival after relapse (19). Conversely, most human sporadic adenomatous polyps typically cluster with CMS2, which was also observed for sporadic rhesus tumor samples. This observation is not entirely consistent with results for human CRC, but could reflect that the CMS classifier has been optimized to characterize human tumors and would require some degree of optimization in rhesus samples.

GSEA indicated activation of key pathways—namely cancer stem cell (CSC) signatures and crypt base— in sporadic MSI rhesus CRC, which corroborates a previously described signature of human MMRd CRC (20). The pathway enrichment between MSI-L/MSS and MSI-H indicates that these advanced, late-stage lesions are transcriptomically similar, which may be driven by the late time point rather than MSI status. These findings provide strong evidence to support the use of these rhesus macaques as a superior animal model for understanding the molecular basis and TME of MSI CRC tumorigenesis.

To quantify the mutational rate in rhesus MMRd CRCs, we leveraged RNAseq data of rhesus LS tissues. We acknowledge that this is not the most optimal way to analyze mutations but allowed us to observe high mutation rates in genes commonly mutated in CRC, independent of MSI status, thus adding additional support to the case for utilization of rhesus macaques for vaccine research, immunotherapy development, and biomarker studies for early detection screening.

We acknowledge that this study has several limitations necessitating further investigation. Importantly, the comparator group, MMR proficient (MMRp) tumors, only included one rhesus, which challenged the validity of the comparison between MMR proficiency and deficiency. Thus, a stronger comparator group is necessary to strengthen our findings. Furthermore, this study lacks pertinent information regarding the timeline of carcinogenesis for both sporadic and LS rhesus tumors, which restricts our understanding of pre-cancer biology, and the timing of tumor development and evolution. Additionally, neoantigen detection and T-cell receptor (TCR) profiling would be an important asset for a complete understanding of the immune system in rhesus macaque CRC. Lastly, our mutation calling was performed using total RNA sequencing data, which although adequate, is less ideal than whole exome sequencing.

In conclusion, this study provides a robust molecular and genetic characterization of a spontaneous and translationally relevant NHP animal model useful for understanding MMRd CRC, including LS CRC.

These results justify the preclinical use of rhesus to study LS CRC and the larger group of sporadic MSI CRCs. Unlike well-established murine animal models and *ex-vivo* cultures, the rhesus MMRd model presented in this study, which occurs in an outbred species with inter-individual variation more representative of the human condition than laboratory mice, affords the ability to test CRC prevention strategies, assess TME dynamics, develop treatment modalities, and survey the immune landscape.

## Material and Methods

### Animal care

The rhesus macaque colony detailed in this manuscript was housed and maintained at MDACC KCCMR in Bastrop, TX. The breeding colony of Indian-origin rhesus macaques (*Macaca mulatta*) at KCCMR is a closed breeding colony, which is specific pathogen free (SPF) for Macacine herpesvirus-1 (Herpes B), Simian retroviruses (SRV-1, SRV-2, SIV, and STLV-1), and *Mycobacterium tuberculosis* complex. All animal experiments were approved by the institutional animal care and use committee (IACUC) and the care of the animals was in accordance with institutional guidelines (IACUC protocol #0804-RN02). Animal care and husbandry conformed to practices established by the Association for the Assessment and Accreditation of Laboratory Animal Care (AAALAC), The Guide for the Care and Use of Laboratory Animals, and the Animal Welfare Act. Tissue specimens from the proximal colon (n=20), the ileocecal junction (n=16), cecocolic junction (n=2), cecum (n=2), and jejunum (n=1), as well as blood samples of rhesus macaques, were collected opportunistically at necropsy following euthanasia for clinical reasons. Formalin-fixed paraffin-embedded (FFPE) blocks and hematoxylin and eosin (H&E) slides were prepared by veterinary pathology technicians and the diagnosis confirmed by veterinary (C.L.H.) and human pathologists (M.W.T).

### Nucleic acid extraction

Macro-dissection was performed to decrease the admixture of adjacent normal tissue and to enrich the percentage of tumor material for subsequent DNA and RNA extraction. De-paraffinization of FFPE tumor and adjacent normal specimens was performed using QIAGEN de-paraffinization solution (QIAGEN, Valencia, CA). DNA and RNA from 19 tumor and adjacent normal samples was extracted using the AllPrep DNA/RNA FFPE Kit (QIAGEN) following the manufacturer’s protocol. In the case of the unavailability of FFPE samples, genomic DNA and RNA were extracted from fresh frozen tumor (n=2) and normal (n=3) samples using the ZR-Duet DNA/RNA MiniPrep extraction kit (ZYMO RESEARCH, Irvine, CA). Quantification was performed with a NanoDrop One™ spectrophotometer (Thermo Fisher Scientific, Waltham, MA) and Qubit™ Fluorometer 2.0 (Qubit, San Francisco, CA) using dsDNA and RNA assay kits. RNA integrity was analyzed using the Tape Station RNA assay kit (Agilent Technologies, Santa Clara, CA).

### Panel design for MSI testing

Commonly used human MSI markers (BAT25, BAT26, BAT40, D10S197, D18S58, D2S123, D17S250, D5S346, β-catenin, and TGFβRII) were used as a reference to design a panel of rhesus MSI markers (21, 22). In brief, genomic positions of human MSI markers in the rhesus macaque genome (rheMac8) were identified using the batch coordinate conversion tool (liftOver) in the UCSC genome browser (23). Repeat patterns were compared to human MSI markers (**Table S1**). Orthologous microsatellite regions corresponding to human MSI markers D2S123, D17S250, and D5S346 were not specific to assess MSI in the rhesus genome. Therefore, they were excluded from the final MSI rhesus panel. Primer sequences to target identified microsatellite regions in rhesus were designed using the NCBI Primer Blast tool (Accession ID# GCF_000772875.2) (24). The primer efficiency was evaluated using the UCSC Genome Browser In-Silico PCR tool (23) with rheMac8 as a reference control. The Baylor College of Medicine genome database was used to calculate the probability of encountering SNPs within the primer sequences. Primers sequences with allele frequency greater than 0.05% were redesigned (**Table S2**).

### PCR-based MSI testing in rhesus CRC

Multiplex PCRs were designed with at least 25 bp size differences among PCR amplicons to afford clear distinction and identification on electropherograms from the Agilent Bioanalyzer 2100. All markers were amplified in 25 μl PCR reactions using 12.5 μl of AmpliTaq Gold™ 360 PCR master mix (Thermo Fisher Scientific, Waltham, MA), corresponding primer sets, and 10 ng of FFPE DNA. Multiplex PCRs were performed in a Veriti 96 Well Thermal Cycler (Applied Biosystems®, Foster City, CA) under the following cycling conditions: initial denaturation at 95°C for 10 min, followed by 35 cycles at 95°C for 30 sec, 55°C for 30 sec, and 72°C for 30 sec. A final extension at 70°C for 30 min was implemented to aid non-template adenine addition. Multiplex PCR products were resolved on a 5% ethidium-bromide stained agarose gel. Multiplex PCRs were analyzed via Agilent 2100 Bioanalyzer DNA 1000 kit (Agilent Technologies, Santa Clara, CA). Electropherograms of adjacent normal and tumor tissue samples were compared to assess the status for each of the MSI markers. Per NCI recommendations, MSI status was assigned by counting the number of unstable MSI markers and samples were assigned to either: MSS (stable markers), MSI-L (1 unstable marker, ≤ 30 %), or MSI-H (2 or more unstable markers, (≥ 30 %) (13).

### MSI testing via fragment analysis for validation of the RheBAT26 and RheD18S58 markers

Fragment analysis (Applied Biosystems®, Foster City, CA) was performed to validate MSI results from the Agilent 2100 Bioanalyzer for RheBAT26 and RheD18S58 MSI markers. In brief, the 5’ end of the forward primer sequences for RheBAT26 and RheD18S58 MSI markers was labeled with a 6-FAM fluorescent dye (Thermo Fisher Scientific, Waltham, MA). A multiplex PCR was designed to amplify RheBAT26 and RheD18S58 MSI markers with labeled primer sequences. PCR master mix and conditions were adopted from well-established PCR experiments. The fragment analysis method was performed by the Advanced Technology Genomics Core at MDACC.

### Sanger sequencing for discovery of germline MLH1 and somatic BRAF mutations

Primer sequences were designed to target *de novo* stop codon *MLH1* and *BRAF* mutations following previously described procedures (see panel design section, **Table S2**). PCRs were performed using the Veriti 96 Well Thermal Cycler (Applied Biosystems®, Foster City, CA) under the following cycling conditions: initial denaturation at 95°C for 10 min, followed by 35 cycles at 95°C for 30 sec, 55°C for 30 sec and 72°C for 30 sec, with a final extension at 72°C for 7 min. Purification of PCR products was performed with an in-house ExoSAP solution [50 μl of Exonuclease I (20,000 units/ml) (NEB® M0568, Ipswich, MA); 40 μl of Antarctic Phosphatase (5,000 units/ml); 16 μl of Antarctic Phosphatase buffer (NEB® M0289S, Ipswich, MA); 144 μl of nuclease-free H2O]. PCR conditions for purification of PCR products were incubation at 37°C for 15 min and at 80°C for 15 min. Quality control of PCR products and purified PCR products was performed running 1% Agarose gel prepared with 25 ml of 1X TBE buffer and 1.2 μl of EtBr. Then, gel-purified PCR products were sequenced by the MDACC sequencing core (ATGC) via the Sanger Sequencing method. Analysis of Sanger sequencing data was performed using DNASTAR lasergene software.

### Immunohistochemistry (IHC)

Immunohistochemistry (IHC) staining for MLH1, MSH2, MSH6, and PMS2 was performed in FFPE tissue sections. Tissue sections were cut at 4 µm and submitted to the MDACC Research Histology, Pathology, and Imaging Core (RHPI) in Smithville, TX. The following Agilent Dako IHC antibodies were used according to manufacturer’s recommendations: IR079, Monoclonal Mouse Anti-human Mutl Protein Homolog 1, clone ES05 for MLH1; IR085, Monoclonal Mouse Anti-human Muts Protein Homolog 2, clone FE11 for MSH2; IR086 Monoclonal Rabbit Anti-human Muts Protein Homolog 6, clone EP49 for MSH6; IR087, Monoclonal Rabbit Anti-Human Posteiotic Segregation Increase 2, clone EPS1 for PMS2 (10).

### Total RNA Sequencing

Truseq stranded total RNA library preparation kit (Illumina®, San Diego, CA) was used to prepare libraries of 21 tumors and 20 matched normal RNA samples, which were extracted from FFPE and frozen tissue samples. Prepared libraries were sequenced for 76nt paired-end sequencing on HiSeq™ 4000 and NovaSeq6000™ sequencers (Illumina®, San Diego, CA).

### Assessment of DNA methylation testing of MLH1

DNA methylation analysis of the *MLH1* gene was performed on DNA from frozen tissue samples of 7 tumors and 3 normal tissue (duodenum and blood) samples using a targeted NGS assay (EpigenDx, Hopkinton, MA). In brief, the bisulfite-treated DNA samples were used as a template for PCR to amplify a short amplicon of 300-500 bp using a set of primers that cover the *MLH1* genomic sequence at −4 kb to + 1kb from the transcriptional start site (TSS). Later, methylation libraries were constructed for methylation analysis on the Ion Torrent instrument at EpigenDx.

### DNA methylation assessment via reduced representation bisulfite sequencing (RRBS)

DNA libraries of RRBS were constructed from FFPE tissue samples of 14 tumors/adjacent normal tissue pairs using the Ovation RRBS Methyl-Seq System at The Epigenomics Profiling Core (EpiCore) of MDACC. In preparation, DNA was digested with a restriction enzyme and selected for size based on established protocols used in the EpiCore. Post-adapter ligation ensured enrichment for CpG islands, and DNA was bisulfite-treated, amplified with universal primers, and qualified libraries were then sequenced on Novaseq6000™ and MiSeq sequencers at the UTMDACC ATGC.

### Bioinformatics Analysis

The FASTQC toolkit was performed for quality control of FASTQ files generated from RNA sequencing (25). The fastp tool was performed to trim adapters and low-quality reads (26). Fasta and gtf files of the reference genome (Mmul_8.0.1) were downloaded from the Ensembl genome browser (27). The reference genome was indexed using the STAR RNA sequencing aligner. Cleaned reads of total RNA sequencing were aligned to the reference genome using the STAR RNA sequencing aligner. Gene level estimated read counts were calculated by STAR RNA sequencing aligner and were saved in reads per gene tabular files (28). This pipeline was implemented on the high-performance computing (HPC) cluster of MDACC. As performed for total RNA sequencing, RRBS FASTQ files were quality controlled using the FASTQC toolkit (25). TrimGalore was performed to trim adapters and low-quality reads. Diversity trimming and filtering were completed with NuGEN’s diversity trimming scripts. Processed fastq files were aligned to the reference genome (Mmul_10) with bismark bisulfite mapper. The methylation information was extracted with bismark methylation extractor script.

Gene expression analysis of RNA sequencing samples with less than 50% uniquely mapped alignment scores were excluded from downstream analyses. Count data per each sample generated by STAR RNA sequencing aligner was combined into one matrix for downstream bioinformatics analyses. Genes that have more than a sum of 100 reads in all samples were excluded from the analysis. The estimated read counts of samples were normalized with variance stabilizing transformation (VST) using the DESeq2 Bioconductor R package (21, 29-31). MSI-L and MSS CRC cases were combined together based on previous human studies. Significant differentially expressed genes between MSI-H and MSS/MSI-L rhesus CRC were calculated using Benjamini-Hochberg (BH)-adjusted P-value ≤ 0.05 and log2 fold change ≥-1 and log2 fold change ≤1. Unsupervised hierarchical clustering was performed via Pearson’s correlation. Comparisons of MMR gene counts between tumor and adjacent normal colorectal mucosa were performed using the DESeq2 Bioconductor R package. Complex heatmap and an enhanced volcano plot were created in R studio (version 3.6.1) (32). Rhesus Ensembl gene-IDs were converted to human Entrez ID for the CMS classification and GSEA. CMS classification of tumor samples was predicted using the random forest (RF) predictor in CMSclassifier R package (version 3.6.1) (14, 19). CMS classification was assigned to the subtype with the highest posterior probability. GSEA was performed with 1,000 permutations using CRC pathways with the fgsea R package (14, 33). CRC pathways included signatures of interest in CRC, the ESTIMATE algorithm that assesses immune and stromal cell admixture in tumor samples, canonical pathways, immune signatures, and metabolic pathways (33, 34).

Somatic and germline variant analyses of rhesus CRC samples were performed following GATK best practices. Filtered variants by Mutect2 and Haplotypecaller tools of GATK were annotated with Variant Effect Predictor (VEP) (35). Mutation rates were calculated by dividing the number of non-synonymous somatic mutations by the number of callable bases.

Species comparison using TCGA datasets utilized raw RNA-Seq counts of MSI-H and MSS colorectal tumor samples and corresponding normal tissue samples (the 2016-01-28 analyses) of the TCGA project COADREAD and MSI status information was downloaded via FirebrowseR (version 1.1.35) package (36, 37). Then the raw data was filtered (min.count = 10, min.total.count = 15, large.n = 10, min.prop = 0.7) and normalized (TMM method) by package edgeR (version 3.32.1) (38). Genes showing statistically significant (BH-adjusted p-value < 0.05) changes in the expression level by at least two-fold (log2FC =1) between MSI-H and MSS samples were identified for the following analysis. The rhesus homologs were found by the Ensembl genome database via the biomaRt package (version 2.46.3) (39-41). Mean CPM (counts per million) of each in COADREAD MSI-H tumor tissues, COADREAD MSS tumor tissues, COADREAD normal tissues, rhesus LS tumor tissues, and rhesus LS normal tissues were used to calculate the Pearson’s correlation of each group. CPM of each gene was used to perform the unsupervised hierarchical clustering, and to generate the dendrogram tree and heat map for individual samples.

For DNA methylation analysis of RRBS, PCA and sample clustering were performed using cytosine report files in methylKit Bioconductor R package (35). The minimum coverage depth was 10 reads. Differentially methylated regions (DMR) were calculated using bismark coverage report files with edgeR Bioconductor R package (26). Significant DMRs at CpG loci were displayed at an FDR of 5%.

## Abbreviations

CRC: colorectal cancer
CSC: cancer stem cell
COAD: Colorectal adenocarcinoma
DEGs: differentially expressed genes
FAP: familial adenomatous polyposis
GSEA: gene set enrichment analysis
H&E: hematoxylin and eosin
Het: heterozygous
IHC: immunohistochemistry
LS: Lynch Syndrome
MMRd: mismatch repair-deficient
MMRp: MMR-proficient
NES: normalized enrichment score
RNAseq: RNA sequencing
MDACC: The University of Texas MD Anderson Cancer Center
TCGA: The Cancer Genome Atlas
READ: rectal adenocarcinoma

## Author’s contributions

EV, SBG, JR, KMS conceived and supervised the study, and provided critical resources to perform the experiments, and wrote the manuscript; NOL designed, performed the experiments, analyzed data, and wrote the manuscript; NOL, RAH, MR and ND performed the analysis of RNA-sequencing data and other bioinformatics analysis; CLH and MWT interpreted pathology slides; SBG and BKD provided the animal model and specimens for analysis; FB genotyped the animals; CMB, LR-U, and KCM provided assistance on the analysis and interpretation of the data, and writing and editorial assistance. All authors critically read and intellectually contributed to the manuscript.

## Acknowledgements

We acknowledge the support of Dr. Awdhesh Kalia at the School of Health Professions of MDACC for providing access to the Agilent 2100 Bioanalyzer for MSI testing analysis. We acknowledge the support of the Advanced Technology Genomics Core (ATGC) for performing the RNAseq, Sanger sequencing, fragment analysis, and RRBS of this project; and Dr. Marcos R. Estecio for RRBS library preparation; and support of the High-Performance Computing facility, which provided computational resources.

