## Supplemental Data for "Comparative Molecular Genomic Analyses of a Spontaneous Rhesus Macaque Model of Mismatch Repair-Deficient Colorectal Cancer"

**Supplemental Figure 1**

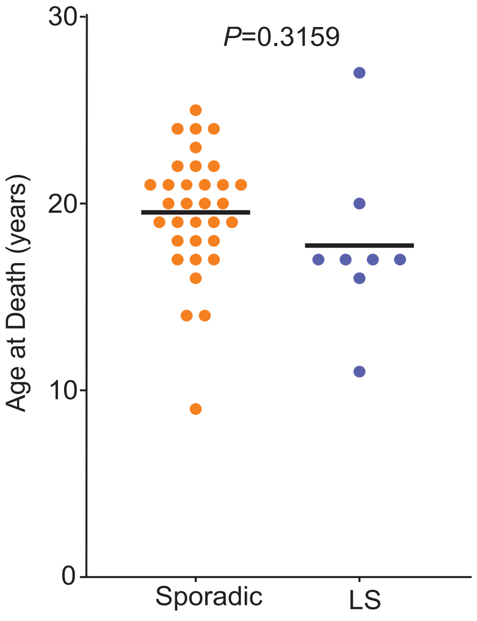


**Supplementary Figure 1. Comparison of mean age at death of rhesus presenting with CRC.** Sporadic animals did not carry a germline mutation in *MLH1*. Mean age of CRC death among rhesus Lynch was younger (17.75 years) compared to the mean age in sporadic rhesus (19.75 years). Welch's t-test, *P*-value=0.3159.

**Supplemental Figure 2**

Healthy rhesus *MLH1*

RM09 *MLH1*

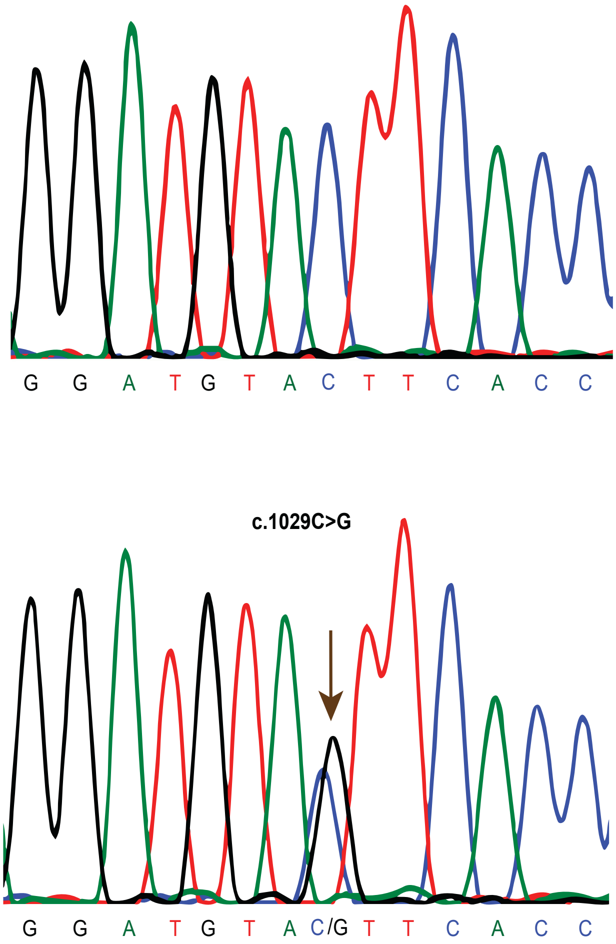


**Supplementary Figure 2. *MLH1* germline mutations detected in rhesus CRC.** Each colored line represents a different type of nucleotide. Brown arrowhead points to the germline mutation detected in normal tissue of rhesus RM09. Non-syndromic animal DNA carries a cytosine (C) nucleotide in c.1029 position of *MLH1*. However, rhesus RM09 carries a mutation in one allele involving the substitution of C>G in c.1029, thus creating a nonsense mutation that leads to a stop codon (TAG).

### Supplemental Figure 3

**A**

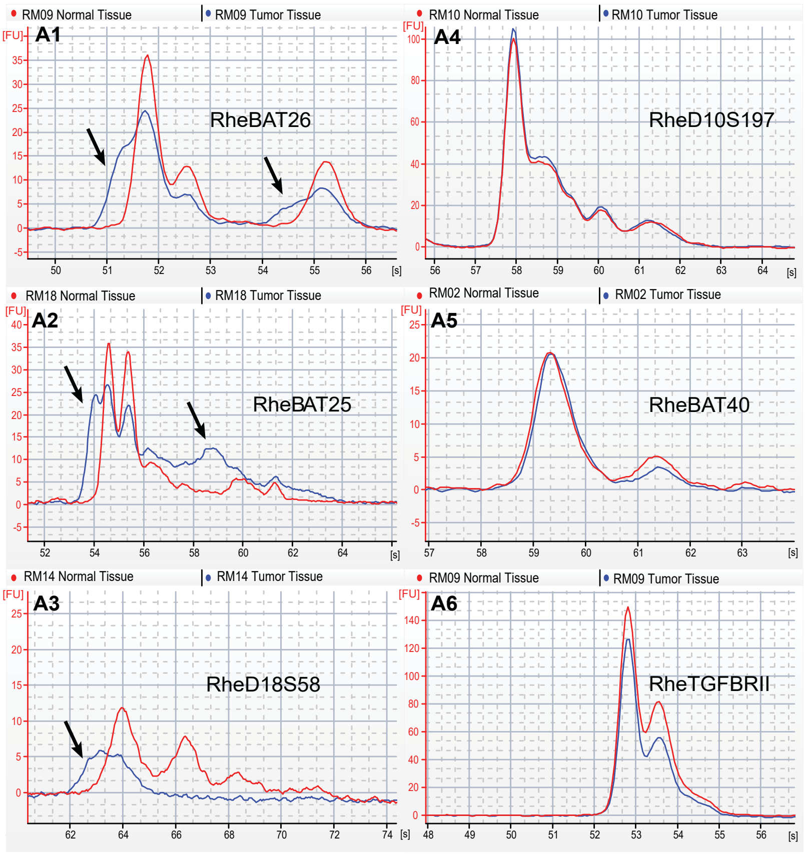


**B**

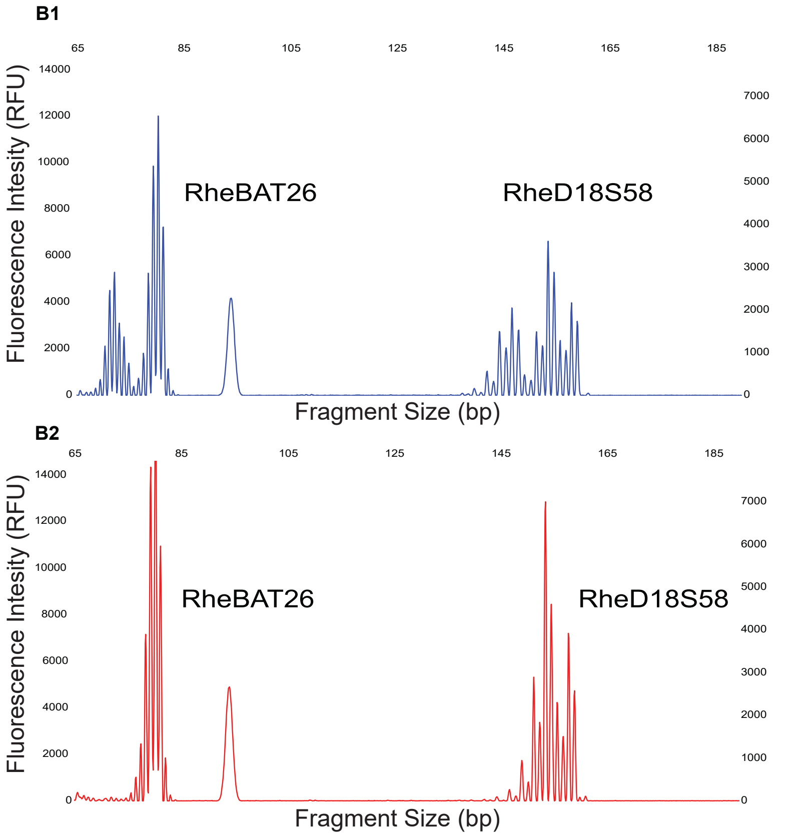


**Supplementary Figure 3. Microsatellite instability analysis.** (A) Examples of microsatellite loci analyzed using the Agilent 2100 Bioanalyzer. Blue and red lines represent tumor tissue and normal tissue, respectively. Electropherograms A1, A2, and A3 are examples of the most frequent microsatellite markers displaying instability in the tumor samples. Arrowheads indicate instability in MSI markers. Electropherograms A4, A5, and A6 are examples of the most common microsatellite markers displaying stability in tumor samples; (B) Examples of microsatellite loci analyzed using fragment analyzer. B1 was generated from tumor tissue of case RM13, displayed instability in markers RheBAT26 and RheD18S58. Arrowheads indicate unstable markers in tumor tissue. B2 was generated from normal tissue of case RM13 and served as a control/reference to establish calls in microsatellite markers in matched tumor tissue of the same case. Overall, fragment analysis validated the results obtained from Agilent 2100 Bioanalyzer calls in markers RheBAT26 and RheD18S58.

### Supplemental Figure 4

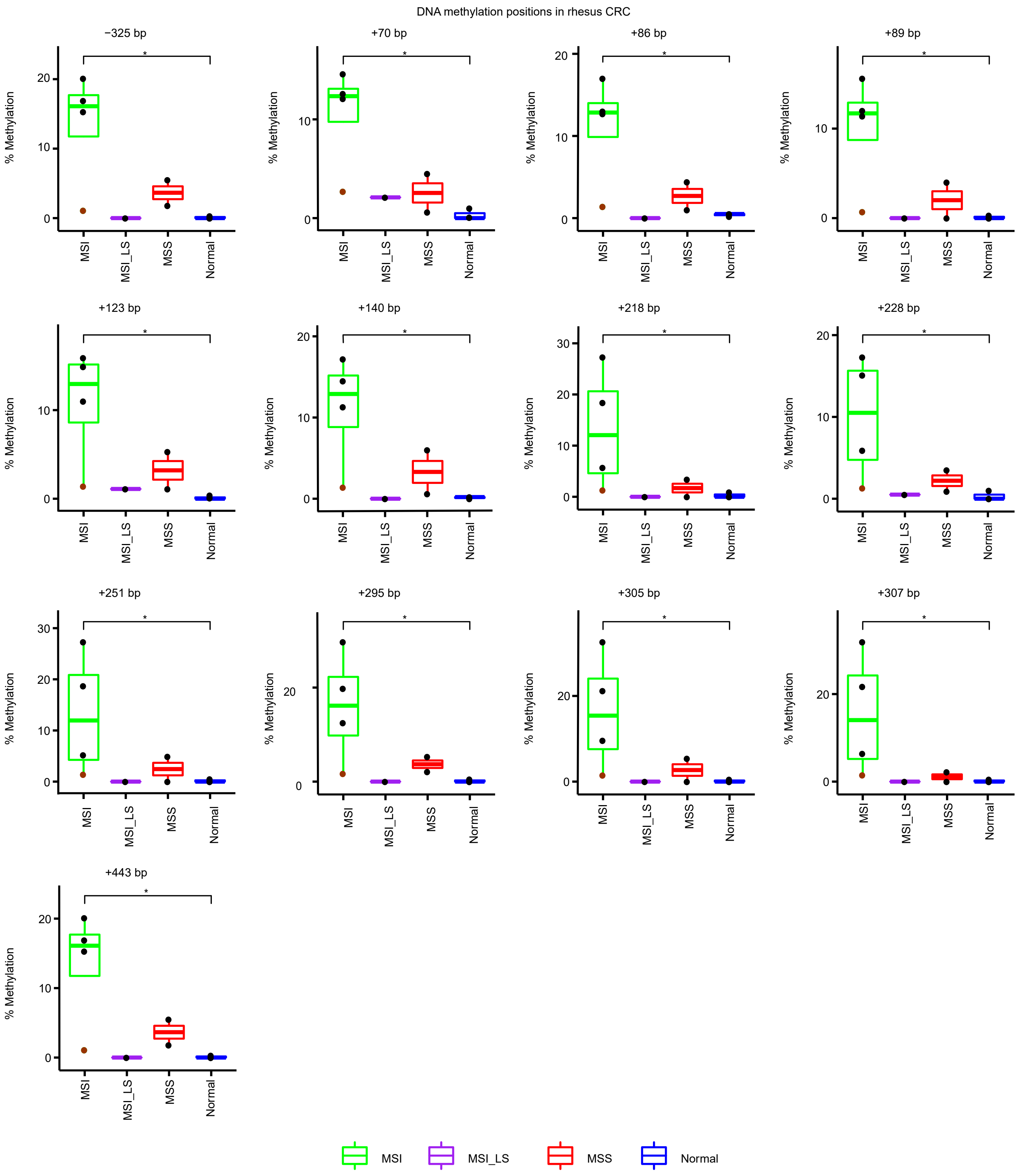


**Supplementary Figure 4. DNA methylation in the promoter region of *MLH1* in rhesus CRC.** The location of CpG islands are shown from the TSS of *MLH1*. A total of thirteen CpG regions were significantly methylated in sporadic MSI-H rhesus CRC samples (**P*-value<0.05). The majority of methylated CpG regions were located in exon 1 of *MLH1*. There were no significant differences in methylation between tumor and normal samples.

**Supplemental Figure 5**

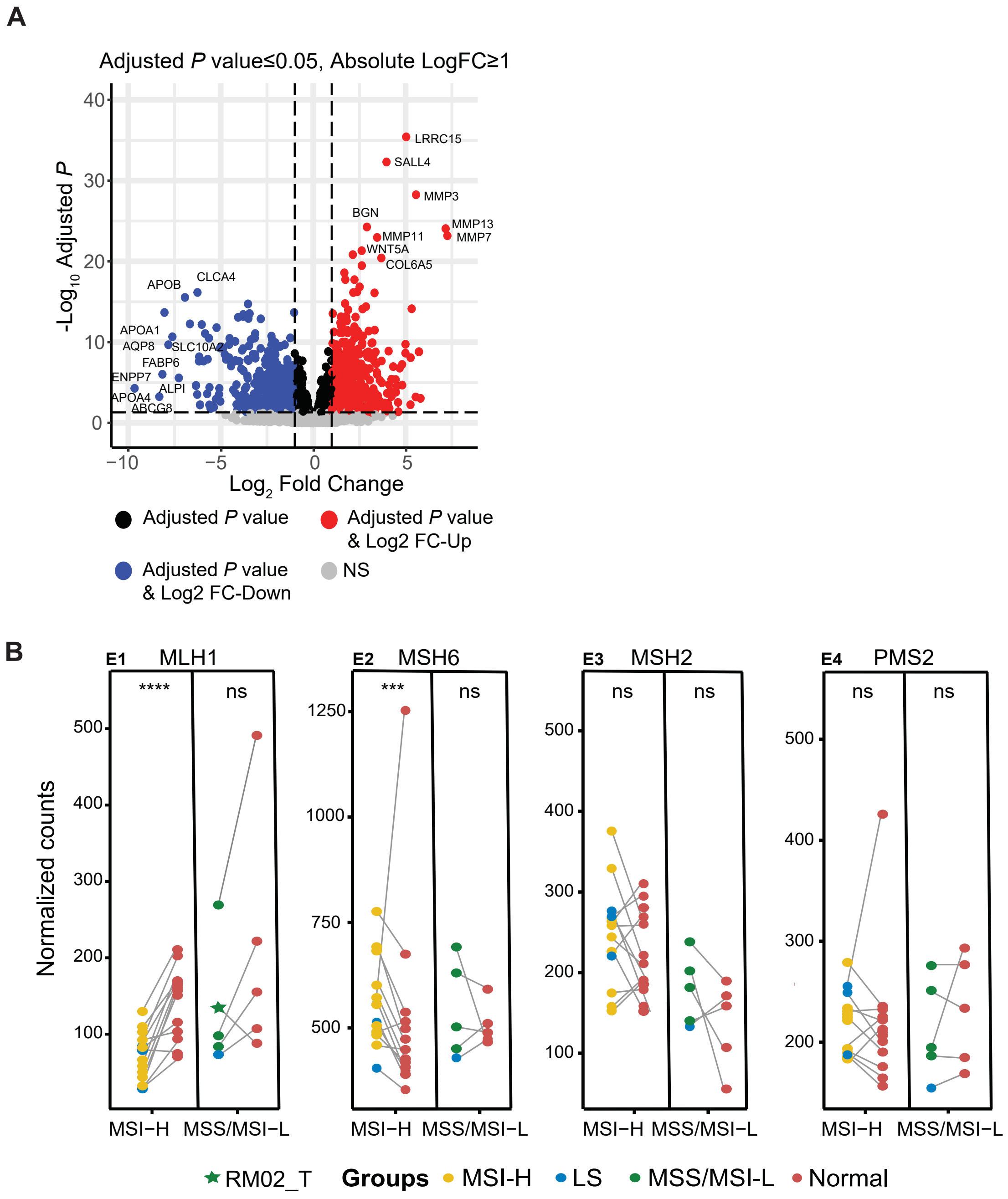


**Supplementary Figure 5. Expression data of rhesus CRC.** (A) Differentially expressed genes between rhesus colorectal normal and tumor samples. Gene expression is displayed in volcano plots with log2(FoldChange) on the X-axis and -log10(BH-adjusted *P*-value) on the Y-axis. The horizontal dash line represents BH-adjusted *P*-value=0.05. The left and right vertical lines represent log2(Fold-Change)= ±1. Significant genes are labeled as upregulated (red) and downregulated (blue) genes. Some significant genes are annotated; (B) Expression of MMR genes in rhesus CRC. Normalized gene counts of whole transcriptome sequencing with variance stabilizing transformation (VST) are on the Y-axis to display gene expression differences of *MLH1*, *MSH6*, *MSH2*, and *PMS2* genes between matched tumor and normal tissue samples. *MLH1* gene expression was significantly (****$P$-value<0.0001) low in MSI-H tumor tissue samples, while *MSH6* gene expression was significantly (***$P$-value<0.001) higher in MSI-H tumors compared to matched adjacent normal. RM02_T (green star) is the only CRC case with a higher expression of *MLH1* in tumors compared to the matched adjacent tissue sample.

**Supplementary Figure 6**

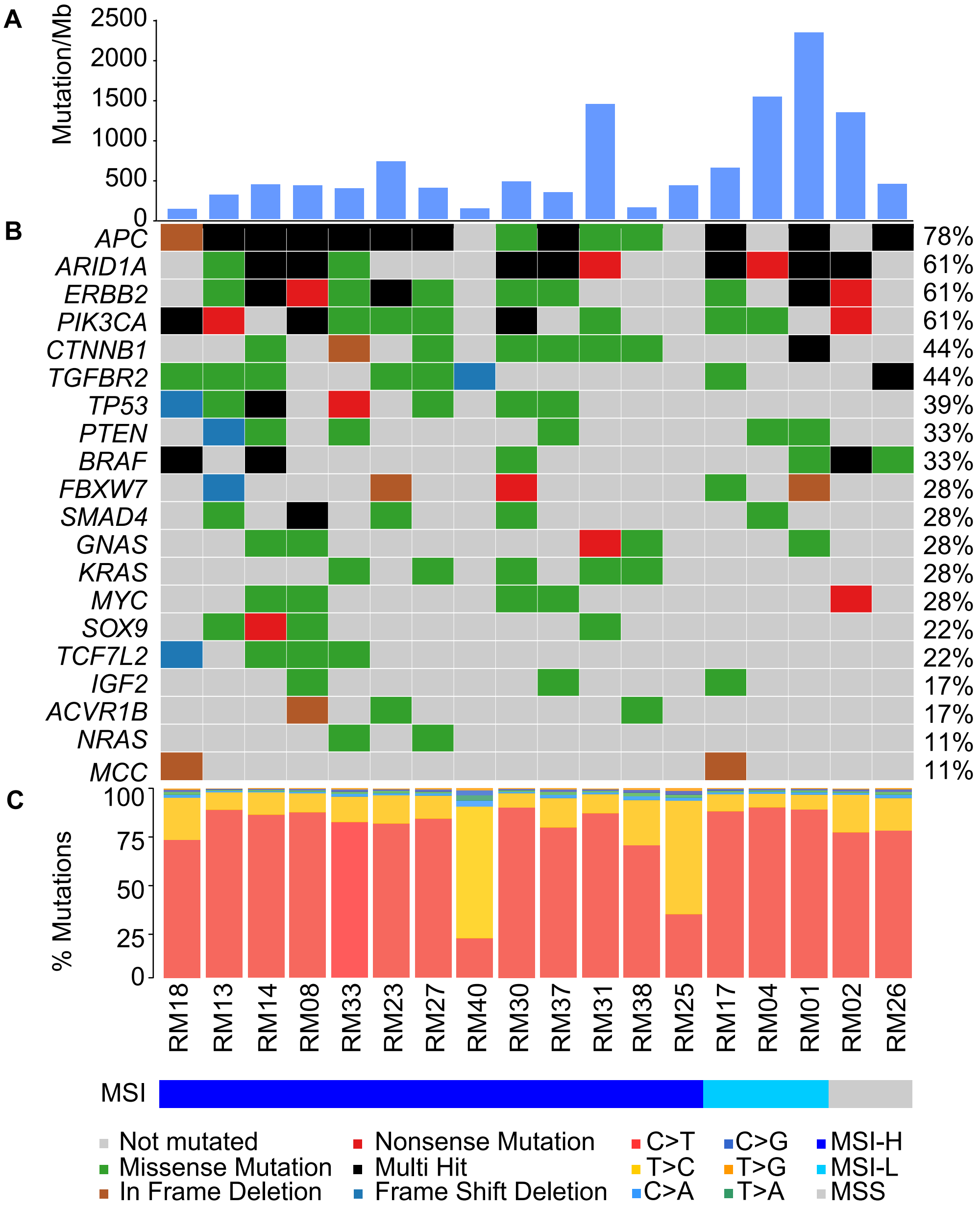


**Supplementary Figure 6. Analysis of somatic variants in rhesus CRC.** (A) The non-synonymous mutation rate is expressed as mutations per megabase (Mb); (B) Commonly mutated genes in human CRC are also altered in rhesus CRC. Each color represents different somatic variants as labeled at the bottom. Black displays multi-hit variants when the sample has more than one somatic alteration in that gene; (C) Proportions of base-pair substitutions in somatic variants in rhesus CRC. Each color demonstrates a different substitution type with C>T being the most abundant in rhesus CRC.

**Supplemental Figure 7**

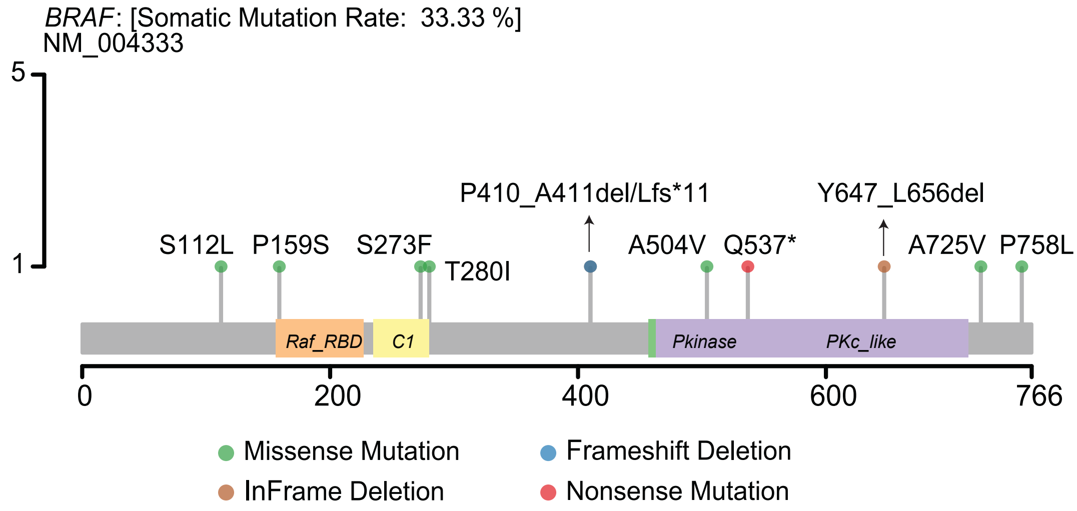


**Supplementary Figure 7. Somatic mutations in *BRAF*.** Missense, nonsense, in-frame deletion, and frameshift deletion mutations detected in BRAF. No hotspots were detected.

**Supplementary Table 1.** Comparison of human and rhesus MSI markers.

| | Repeat Patterns | Rhesus MSI Markers | Repeat Patterns |
|---|---|---|---|
| **BAT 25** | $(A)_{25}$ | c-kitRheBAT25 | $(A)_{36}$ |
| **BAT 26** | $(A)_{26}$ | RheBAT26 | $(A)_{27}$ |
| **BAT 40** | $(T)_{7}$….$(T)_{40}$ | RheBAT40 | $(T)_{6}C(T)_{6}C(T)_{5}C(T)_{5}C(T)_{4}C(T)_{17}$ |
| **D10S197** | $(CA)_{7}$…$(CA)_{12}$ | RheD10S197 | $(CA)_{18}$ |
| **D18S58** | $(GC)_{5}GA(CA)_{17}$ | RheD18S58 | $(CA)_{18}$ |
| **TGFβRII** | $(A)_{10}$ | RheTGFβRII | $(A)_{10}$ |

**Supplementary Table 2.** Primer sequences for determination of rhesus MSI status and *MLH1* germline mutation.

| | FW Primer (5'-3') | RV Primer (5'-3') | PCR Product (bp) | |
|---|---|---|---|---|
| **c-kitRheBAT25** | CGAGATTGTAC CACTGCAC | TGCCTGGCTGA TATTTCTTTA | 119 | *c-kit* |
| **RheBAT26** | GATATTGCAGCA GTCAGAGC | AACCATTCAACA TTTTTAACCC | 87 | *MSH2* |
| **RheBAT40** | CCTACACCACAA TCCTGCT | GGGTGGTAGAGC AAGACCT | 144 | *3β-HSD* |
| **RheD10S197** | CTTCAGGGTGAAA GGACGAG | ACCTCGAGTGGCA TTTTGAA | 128 | *GAD2* |
| **RheD18S58** | TCCCTTAGGAGGC AGGAAAT | TCCTGGCCGGCTT TATTTAT | 153 | *DCC* |
| **RheTGFβRII** | TGACTTTATTCTGG AAGATGCTG | AACACATGAAGAA AGTCTCACCA | 83 | *TGF-β-RII* |
| **RheMLH1** | TAACAGGCAAAAAT CTGGGC | CCACATACACCATA TGTGCC | 324 | *MLH1* |
| **RheBRAF** | CCTAAAATCTTCAT AATGCTT | ATAGCCTCAATTCT TACCAT | 209 | *BRAF* |

**Supplementary Table 3.** CMS classification of rhesus CRC. This analysis was performed using random forest classifier in the CMSclassifier R studio to establish CMS status in rhesus CRC.

| Sample | CMS1 | CMS2 | CMS3 | CMS4 | Predicted CMS (RF) | MSI status |
|---|---|---|---|---|---|---|
| RM18 | 0.25 | 0.61 | 0.12 | 0.02 | CMS2 | MSI-H |
| RM13 | 0.36 | 0.42 | 0.09 | 0.13 | CMS2 | MSI-H |
| RM14 | 0.2 | 0.4 | 0.13 | 0.27 | CMS2 | MSI-H |
| RM04 | 0.21 | 0.37 | 0.31 | 0.11 | CMS2 | MSI-L |
| RM01 | 0.22 | 0.26 | 0.13 | 0.39 | CMS4 | MSI-L |
| RM02 | 0.12 | 0.6 | 0.17 | 0.11 | CMS2 | MSS |
| RM08 | 0.33 | 0.39 | 0.24 | 0.04 | CMS2 | MSI-H |
| RM17 | 0.14 | 0.12 | 0.03 | 0.71 | CMS4 | MSI-L |
| RM09 | 0.36 | 0.32 | 0.26 | 0.06 | CMS1 | MSI-H |
| RM26 | 0.25 | 0.55 | 0.18 | 0.02 | CMS2 | MSS |
| RM33 | 0.42 | 0.32 | 0.19 | 0.07 | CMS1 | MSI-H |
| RM23 | 0.35 | 0.31 | 0.08 | 0.26 | CMS1 | MSI-H |
| RM27 | 0.28 | 0.42 | 0.28 | 0.02 | CMS2 | MSI-H |
| RM40 | 0.14 | 0.22 | 0.1 | 0.54 | CMS4 | MSI-H |
| RM30 | 0.24 | 0.48 | 0.23 | 0.05 | CMS2 | MSI-H |
| RM37 | 0.42 | 0.42 | 0.13 | 0.03 | CMS1-CMS2 | MSI-H |
| RM31 | 0.24 | 0.63 | 0.1 | 0.03 | CMS2 | MSI-H |
| RM38 | 0.44 | 0.22 | 0.21 | 0.13 | CMS1 | MSI-H |
| RM25 | 0.29 | 0.25 | 0.12 | 0.34 | CMS4 | MSI-H |
